# 3D nanoscale analysis of bone healing around degrading Mg implants studied by X-ray scattering tensor tomography

**DOI:** 10.1101/2020.11.09.375253

**Authors:** Marianne Liebi, Viviane Lutz-Bueno, Manuel Guizar-Sicairos, Bernd M. Schönbauer, Johannes Eichler, Elisabeth Martinelli, Jörg F. Löffler, Annelie Weinberg, Helga Lichtenegger, Tilman A. Grünewald

## Abstract

The nanostructural adaptation of bone is crucial for its compatibility with orthopedic implants. The bone’s nanostructure determines its mechanical properties, however little is known about its temporal and spatial adaptation in degrading implants. This study presents insights into this adaptation by applying electron microscopy, elemental analysis, and small-angle X-ray scattering tensor-tomography (SASTT). We extend the SASTT reconstruction to multiple radii of the reciprocal space vector *q*, providing a 3D reciprocal-space map per voxel. Each scattering curve is spatially linked to one voxel in the volume, and properties such as the thickness of the mineral particles are quantified. This reconstruction provides information on nanostructural adaptation during healing around a degrading ZX10 magnesium implant over the course of 18 months, using a sham as control. The nanostructural adaptation process is observed to start with an initially fast interfacial organization towards the implant direction, followed by a substantial reorganization of the volume around the implant, and an adaptation in the later degradation stages. The study sheds light on the complex bone-implant interaction in 3D, allowing a more guided approach towards the design of future implant materials, which are expected to be of great interest for further clinical studies on the bone-implant interaction.

**TOC text and figure:** Degrading Magnesium implants are mechanically and chemically well adapted orthopedic implant materials and ensure a gradual load transfer during bone healing due to their degradation. The impact of the implant degradation on the bone nanostructure is however not fully understood. This study unveils the processes 3D and shows different stages of bone healing.

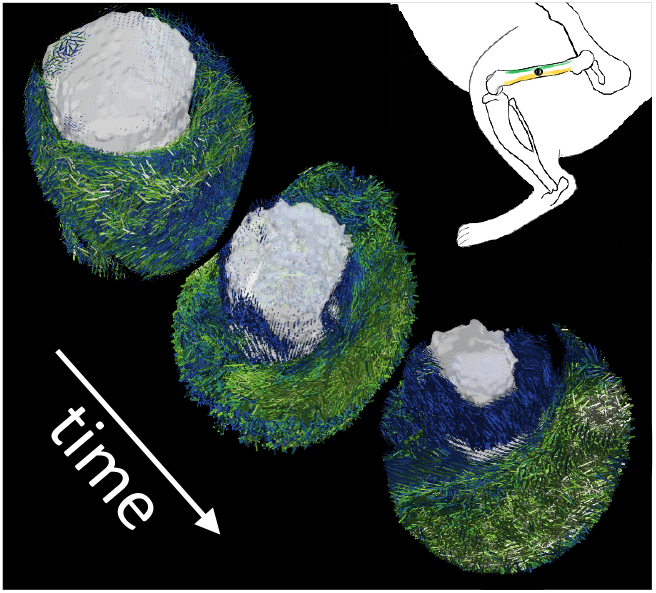

## 1. Introduction

The engineering of orthopaedic implants and their paradigms of usage have seen great change in recent years. The materials employed had properties that spread over bio-inert, biocompatible, and bio-active. The improvements related to this change, however, cannot be reflected solely by modifications to the implant materials, but requires further understanding of the nanostructural adaptation of bone healing with respect to the degradation of such implants. Therefore, there is a great interest in detailed investigation of the adaptation of bone during implant degradation and the corresponding implant-bone interaction at the interface.

Bone structure shows remarkable complexity and is organized over 9 orders of magnitude. These orders range from the fractal organization of the mineral particles to the macroscopic bone shape, which is thought to contribute to its mechanical optimization. In long bones, three major constituents can be identified: trabecular bone, cortical bone and the bone marrow. Each of these components shows a rich functional structure on the micro- and macroscale, as well as different organizational motifs on the meso-to nanoscale[1]. Despite the ongoing discussion on the organization of the nanostructure, it is generally agreed that the nanostructured building blocks of bone are collagen fibrils, extrafibrillar matrix proteins and hydroxylapatite (HAP) mineral particles. In the classical model [2], the collagen fibres are are mineralized in the gap zones of the overlapping collagen molecules. The mineral particles inside a single collagen fiber are thought to be layered with respect to each other[3], this layered register even extending over several fibrils (~500 nm) in the case of mineralized turkey leg tendon [4]. Over larger lengths scales of few μms, the individual layers are thought to be rotated around the collagen fiber axis, while retaining their preferred orientation along the collagen fiber axis.[5]

The understanding of bone nanostructure is an active research area. However, only recently research started to focus on the nanostructural adaptation of bone triggered by implant placement [6]. Besides classical techniques such as histology, computed tomography and electron-based methods to investigate the bone structure, small-angle X-ray scattering (SAXS) has become key to uncover the bone nanostructure and its adaptation. Fratzl and coworkers [7] contributed pioneering work on the nanostructural arrangement of mineral particles in bone and the analytical treatment of SAXS data [8]. The interaction between implants and bone has become since a widely investigated research area. Noteworthy publications on nondegrading implants include the work of Besenbacher *et al*. [9], which showed that mineral platelets have orientational rearrangement around a non-degrading titanium (Ti) screw implant. Hoerth *et al*. [10] showed the bone-growth process around zirconium (Zr) and Ti implants. In addition, high-speed X-ray absorption tomography allows new insights into the degradation behaviour of magnesium (Mg) implants by time-resolved imaging [11].

The advent of synchrotron-based scanning-beam techniques provides insights with micrometer resolution, which generates an understanding of complex structural changes that often happen at length scales below 100 μm. By using microbeam scanning diffraction techniques, we previously showed significant alterations in the bone nanostructure within 20 μm from the bone-implant interface. The interface is characterized by high Mg levels and altered mineral particle size and mineral structure [12]. Recent work has also illustrated the adaptation of the bone nanostructure as a response to degrading implants [13]. In that work, a 2D scanning technique was employed and two findings were particularly striking: (i) the early onset of nanostructural reorientation (after 1 month), which, however, showed no evident change in the mineral particle size and (ii) the late onset of bone growth after 12 months and persistent orientation effects after 18 months. These findings were corroborated by mechanical studies that confirmed the slow healing rates, based on the hardness and stiffness of the bone in the vicinity of the degrading implants [14]. The main conclusion of that work was that Mg implants induce a localized adaptation of the nanostructure, which needs to be considered in the design of degrading metallic implants. Nevertheless, it should be noted that the 2D study was naturally limited to scanning the sample in the plane of the cross-section, making it impossible to detect any potentially more intricate orientation patterns, such as a wrapping around the implant. Focusing on the positive side effect of initiating bone growth [15] due to the release of Mg ions and matching this with already reported deployment in orthopaedics [16], potential further applications may be found in Mg-based scaffolds [17, 18] with a focus on nanostructural integration.

The Mg alloy of this study is ZX10 [19], which is a lean Mg alloy containing 1.0 wt.% Zn and 0.3 wt.% Ca. ZX10, is especially attractive for biodegradable implants because it is composed entirely of elements essential for the human body, and is thus intrinsically biosafe. This composition is distinguishable from other Mg alloys that contain rare-earth elements, such as yttrium (Y) [20], which gives rise to some concerns as Y accumulates in bone tissue [21] and might generate an adverse immunological response [22]. ZX10 contains intermetallic precipitates (IMPs) on the nanoscale, which governs its degradation behaviour and leads to improved mechanical properties. Its detailed biodegradation behaviour [23] and its susceptibility to stress-corrosion cracking and corrosion fatigue [24] has been studied *in vitro* in simulated physiological environment and its response to bone has also been documented recently via a long-term *in vivo* study [23].

A drawback of scanning-diffraction techniques is their limitation to 2D slices, which also involves destructive sample preparation. This limitation is especially problematic when the structural adaptations are expected to happen in a volume, as one would expect for degrading implants. 3D diffraction tomography has already given promising insights into nanostructural development [25]. The extension of scattering techniques from 2D to 3D has posed a great challenge and was recently solved for oriented nanostructures with a tensor tomography approach by Liebi *et al* and Schaff *et al*.[26, 27] This development now enables the study of the bone nanostructure [28] or even whole bone-implant structures in 3D and finally allows to address some pressing questions with respect to degradable Mg implants, such as: (i) How does the implant degradation process affects the bone nanostructure in 3D around the implant? And (ii), what is the influence of the degradation products on the bone nanostructure?

Here, we present new insights into the interaction between degrading Mg implants and bone nanostructure. To this end, we investigated the degradation process and the subsequent bone healing for a biomedically relevant ZX10 implant using small-angle X-ray scattering tensor tomography (SASTT). This method yields the full reciprocal space map for every voxel of a three-dimensional sample while retaining high spatial resolution in 3D (see Methods section for a more detailed description). The 3D structure as a function of the distance to the implant-bone interface and the correlation of its changes to the presumed loading pattern and implant direction is an innovative way to investigate such unknown, complex materials. Based on SASTT reconstructions, we spatially resolved a bone-implant interface that is rich of degradation product, giving a distinct fingerprint to the scattering pattern. The extension and distribution of such interface around the implant volume allowed us to evaluate the implants’ capability for osseointegration.

## Experimental Section/Methods

### Samples

High-purity magnesium was alloyed with zinc and calcium to synthesize the alloy Mg-Zn1.0-Ca0.3 (in wt.%), resp. Mg-Zn0.37-Ca0.18 (in at.%). After solution and aging heat-treatments, indirect extrusion was performed at 325°C to generate ZX10 rods of 3 mm diameter. More details on materials synthesis and processing can be found in Refs.[23, 29]. Pins of 1.6 mm diameter and 8 mm length were then machined using polycrystalline diamond tools, taking special care to avoid any kind of surface contamination. The pins were cleaned using ultrasonic waves, air dried in clean-room atmosphere, and packaged airtight. Sterilization was performed by gamma irradiation at 51 kGray. The outer packaging prevented corrosion attack prior to implantation.

The cylindrical implants were inserted transcortically into the femoral bones of 5-weeks old male Sprague-Dawley rats under general anesthesia. These comparatively young rats were chosen to be able to sample the whole implant degradation cycle with the same animal model. The entire surgical procedure, the post-operation treatment and the final euthanization step for explant analysis are described in[20]. The animal experiments were performed according to approved ethical standards and authorized by the Austrian Ministry of Science and Research (accreditation number BMWFW-66.010/0122-WF/V/3b/2014). After explantation, the femoral bones with the overgrown implants were embedded in PMMA resin (‘Technovit 7100’, Heraeus Kulzer, Wehrheim, Germany) and the whole blocks were scanned via μCT to detect the degradation state and implant location. Representative samples from time points 1, 6 and 18 months were selected and milled into cylinders of 3 mm diameter using a Struers Accutom 50 with a diamond polishing wheel. These cylinders were glued onto metal pins and centered on a goniometer head for the SASTT experiments.

### SASTT measurements

The SASTT experiments were carried out at the cSAXS beamline at the Swiss Light Source (SLS) at the Paul Scherrer Institut, Villigen, Switzerland. A schematic overview of the experiment and data processing pipeline is shown in Figure 1. An X-ray beam energy of 18 keV was selected using a Si (111) double crystal monochromator and focused to a spot size of 50 μm by a combination of the vertically focusing monochromator and a horizontally focusing mirror. The sample was mounted on a steel needle and aligned to the center of rotation of a two-rotation-axis setup, which allows for rotation, tilt, and x-y scanning of the sample. The sample was mapped spatially with a resolution of 50 μm and an exposure time of 0.03 s using fly scans in the x-direction. A schematic sketch of the setup is given in Figure 1a. A total of 319 projections were taken at 10 β tilt angles between 0 and 45° and α rotation angles between 0 and 180° (β = 0°), and 0 and 360° for β ≠ 0. The number of rotations per tilt angle was reduced by a factor of cos(β) to provide equal angular sampling at different tilts[30].

**Figure 1.**
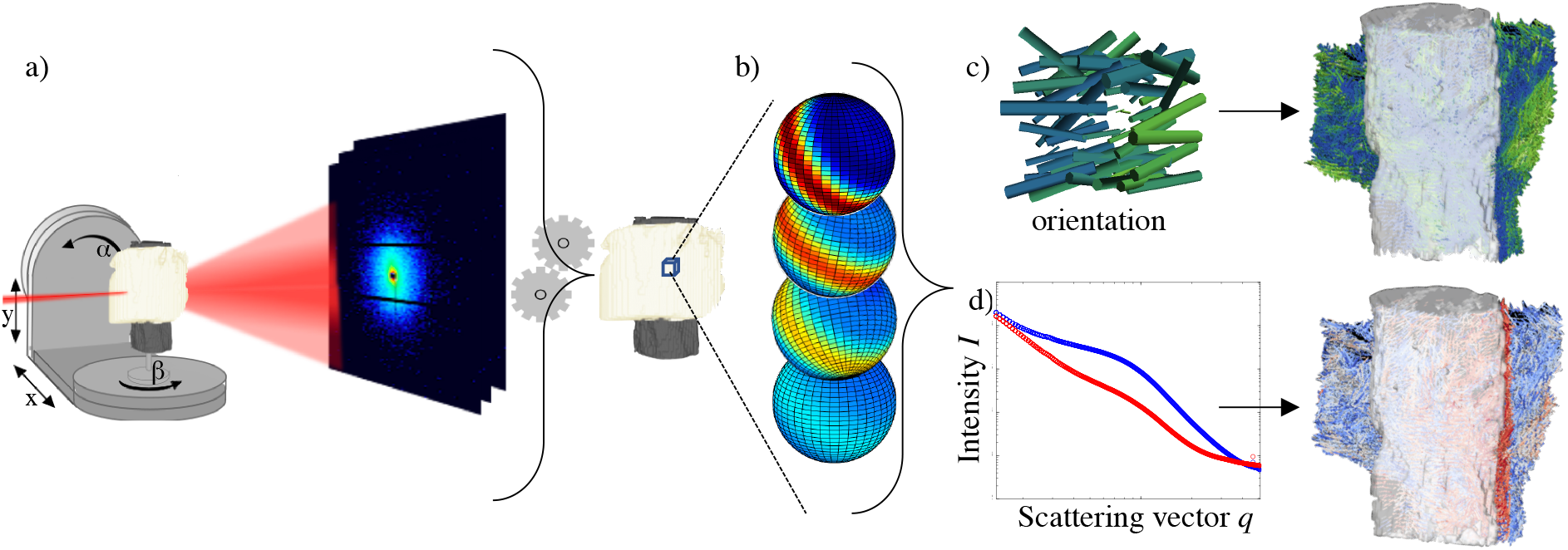
Data acquisition, processing and visualization pipeline. (a) The collection of the SASTT datasets requires a raster scanning of the sample in the x-y direction and the collection of subsequent projections at various angles of rotation β and tilt α. (b-d) After aligning the projections, a 3D reciprocal-space map was reconstructed for every voxel (b) and further analysed with respect to the orientation and DOO (color) (c) or the average scattering curve in every voxel and the low-q exponent (color) (d).

The SAXS patterns were measured using a Pilatus 2M detector[31] with a sample-detector distance of 2 m, which covers a q-range from about 0.1 to 5 nm^-1^, with 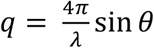, where *λ* is the wavelength of the X-rays, and *θ* is the half scattering angle. A beamstop was used to block the directly transmitted beam. The sample transmission was measured using a photodiode mounted on the beamstop, and this was used to correct the scattering signal for sample transmission[32].

### Reconstruction of SASTT data

The data were integrated by in-house Matlab routines in 2000 radial and 16 azimuthal bins [33]. The alignment of the projections, based on a reference tomogram at α = 0°, and the subsequent SASTT reconstruction were carried out with data in the range of q = 0.43 nm-1 to 0.72 nm-1. The details of the SASTT reconstruction procedure can be found elsewhere[30]. In brief, the reciprocal space map in each voxel is described by a series of spherical harmonics of orders m = [0,0,0,0] and degrees l = [0,2,4,6] and a zenith orientation parameterized with the spherical angles θ and ϕ. Using even numbers for degrees l and zero-order m is based on the expected rotational symmetry around one axis, assuming that the mineral-platelet surface-normal is randomly oriented in the plane perpendicular to the collagen fibril axis, which we can safely assume as we average over millions of mineralized collagen fibrils and about 10^9^ mineral particles in each voxel.[5,26,28,30] These coefficients were optimized using a stepwise approach as further described in Ref.[30]. The validity of the reconstructions was checked by comparing the orientation plots of measured raw data with the reconstructed data. From the full SASTT reconstruction, the direction and DOO of the local nanostructure was extracted, as illustrated in Figure 1c. The DOO, ρ, is thereby defined as

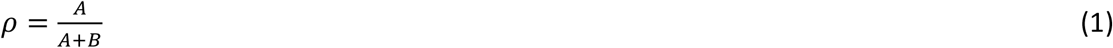

where A is the anisotropic intensity in the spherical harmonics components and B is the isotropic scattering component of the spherical harmonics, in analogy to Pabisch et al.[34] for the case of 2D scattering. For the mathematical expression of ρ in terms of the spherical harmonic coefficients, we refer to Ref.[30]. To highlight the extracted quantities for every voxel, a small sub-volume is shown in Figure 1c, which shows the orientation (as the orientation of the glyph) and the symmetric intensity (in color code).

For the q-resolved reconstruction, only the symmetric scattering intensity was reconstructed with the SASTT code by only optimizing the coefficients for m = 0, l = 0. This optimization was carried out for 200 log-spaced q-bins of the data, resulting in a one-dimensional scattering curve akin to conventional SAXS, as illustrated in Figure 1d. This approach was chosen for computational efficiency as only the averaged intensity of every voxel was used in further analysis. We verified for one sample with reconstructions of the full, higher-order orientation tensor, i.e. orders m = [0,0,0,0] and l = [0,2,4,6] and a zenith orientation parameterized with the spherical angles θ and ϕ, that the obtained angularly averaged intensity was very comparable for each q-bin and did not change the fitted values (see SI Figure 7). This approach is comparable with conventional diffraction CT[25,35–37] or small-angle scattering CT[32] in the sense that it is backprojecting the intensity as a scalar quantity to form a volume, in which every voxel contains a 1D scattering curve. All the projections recorded under all tilt angles α were used to reconstruct the volume, thus it yields the average azimuthal scattering of the 3D reciprocal-space map. In contrast to the SASTT method, in which we need to extract multiple parameters to obtain the orientation tensor, here we just aimed at refining a single scalar quantity in the form of the scattered intensity as a function of the scattering vector *q* (Figure 1d), in order to use established models[34, 38] to fit the scattering curves.

### Fitting of scattering curves

In order to characterize the scattering curves in a more general manner, and to be able to account for the not yet mineralized interface, a power-law fit to the Guinier region was used as a most general descriptor of structure [39]. In brief, it allows to draw a conclusion on the scattering behaviour and the shape of large scattering structures in the system.

The intensity decay takes the form

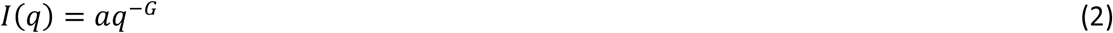

where I(q) is the scattered intensity, A is the amplitude and −G is the exponent. For the power-law decay fit, a q range between 0.3 and 0.7 nm^−1^ was used, as shown in Figure SI 8a. The fitting was implemented with the lmfit python package using a Levenberg-Marquard algorithm. This results in the analysis schematically shown in Figure 1d. With this, one can discriminate between well mineralized bone and the non-mineralized interface region.

In order to evaluate the size of the scattering mineral particles in the mineralized portion of the sample, the azimuthal intensity *I*(*q*) can be evaluated. Assuming a two-phase system with a mineral fraction of 50% [40] and predominantly platelet-shaped particles, the T parameter can be calculated as

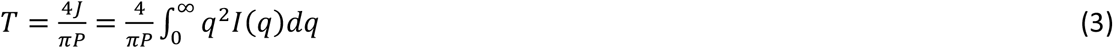

from the Porod constant P and the Invariant J. The Porod constant was determined from the Porod region of the data (q = 0.9 nm-1 to 2.3 nm-1, with the data presented in a Porod plot, Iq^4^ vs q^4^ (see SI Figure 8b), and a linear fit extrapolated to q = 0 to determine the Porod constant (red line in SI Figure 8b). The invariant was determined from the Kratky plot q^2^I vs q (SI Figure 8c) by integrating the curve in the experimentally accessible q-range and extrapolating linearly to q = 0 and with q^-4^ to q = ∞. Determining the parameters in this way follows the approach laid out by Pabisch et al. [34].

In order to evaluate the corresponding parameters as a function of the distance to the implant surface, the absorption tomogram was segmented for air, bone, and implant, and the implant surface was defined by dilating the implant volume and subtracting the original implant volume from it. This operation was followed by a morphological closing operation to yield a closed surface. After this, the distance between each point and the implant was calculated.

### SEM/EDX data acquisition

The SEM/EDX data was acquired with a FEI Quanta 250 FEG. Samples were investigated after the SASTT experiments and to this end re-embedded with an epoxy resin (Agar LV resin) to facilitate handling and cutting with a diamond wheel using a Struers Accutom 50. The sample surface was subsequently polished with 0.1 μm alumina suspension. The samples were at first left unsputtered for BSE and EDX analysis and the SEM was operated in low vacuum mode at 130 Pa. The acceleration voltage was 20 kV. The EDX data was acquired with an EDAX Octan SDD detector with an exposure time of 1000 μs and 100 repetitions.

Subsequently, the sample was sputter-coated with carbon and reinvestigated via SE imaging in high vacuum at an acceleration voltage of 5 kV.

### SEM data treatment

The BSE data were segmented using a trainable machine-learning algorithm (Trainable Weka Segmentation) [41], implemented in the Fiji image processing package. A training dataset was built by a manual selection of the relevant structures (bone, implant, lacunae and background). This segmentation was further used to calculated the lacunar density in Matlab.

### MicroCT data acquisition

μCT data were acquired using a Siemens Inveon Acquisition Workplace 1.2.2.2 in cone-beam geometry. Scans were carried out at 80 kV voltage, 500 μA current and 1050 ms exposure time per projection. A total of 180 projections over 210° were collected. The 3D data were reconstructed using a filtered backprojection algorithm, resulting in a voxel size of 18.9 μm.

## 2. Results

Figure 2 sketches (a) a rat femur with an implant and (b) the orientation of the bone hydroxyapatite mineral particles, where (c) represents the orientation of the samples as shown in Figure 2 (a-c) and Figure 6. As a result of the presumed bending loading mode, zones of tensile and compressive stress are generated on the top and bottom, respectively. Hence, we are able to describe the reaction for the four different quadrants as the distal, proximal, tensile and compressive sides, as shown in Figure 2c. For further discussion we introduce a reference vector along the distal-proximal direction that corresponds to the long axis of the femur (represented by a blue arrow Figure 2c).

**Figure 2.**
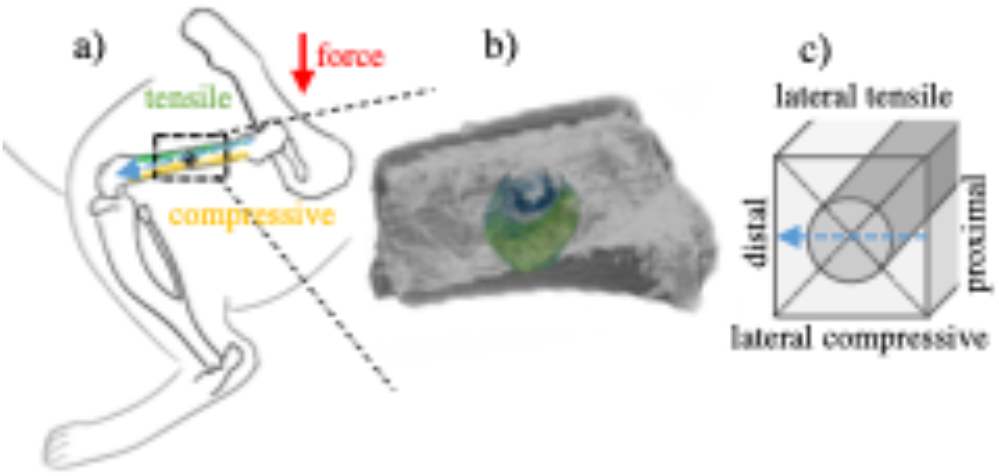
(a) Sketch of implant position and load distribution in a rat bone, with respect to the main bone axis from proximal to distal (blue arrow), (b) μCT measurement overlaid with the SAXS tensor tomography measurements performed, and (c) schematic sketch of the orientations around the implant with the long bone reference vector (blue arrow). The orientation in (c) corresponds to that of Fig. 2 a-c and Fig 5.

Bone samples containing a bioresorbable ZX10 Mg implant in progressing degradation stages (1 month, 6 months and 18 months after implant placement) and control samples, in which a hole was introduced in a similar position but without an implant (sampled 1 month and 6 months after surgery), were investigated by SASTT. The implant/hole placement and the region where the sample has been extracted for SASTT are shown in *ex vivo* μCT scans in SI Figure 1. SASTT gives volumetric nanostructural information on the main scattering direction and degree of orientation (DOO), as well as the scattered intensity in every voxel. The technique and the methodology are further explained in the Methods section. Within each subvolume (i.e. voxel) sampled by SASTT in this experiment (50×50×50 um), we measure the average orientation of several millions of mineralized collagen fibrils and about 10^9^ mineral particles. Due to the rotational averaging of the mineral particles around the collagen fibrils at this length scales[5], there is no clear preferred orientation around the fiber axis, and only the preferred orientation along the fiber axis remains.

The visualization in Figure 3 shows for every voxel of the 3D sample the orientation of the nanostructure (in the direction of the glyph), the scattered intensity (via the length of the glyph) and an additional parameter such as the degree of orientation or the low-*q* exponent, i.e. the Guinier exponent *G*, via the color coding of the glyph.

**Figure 3.**
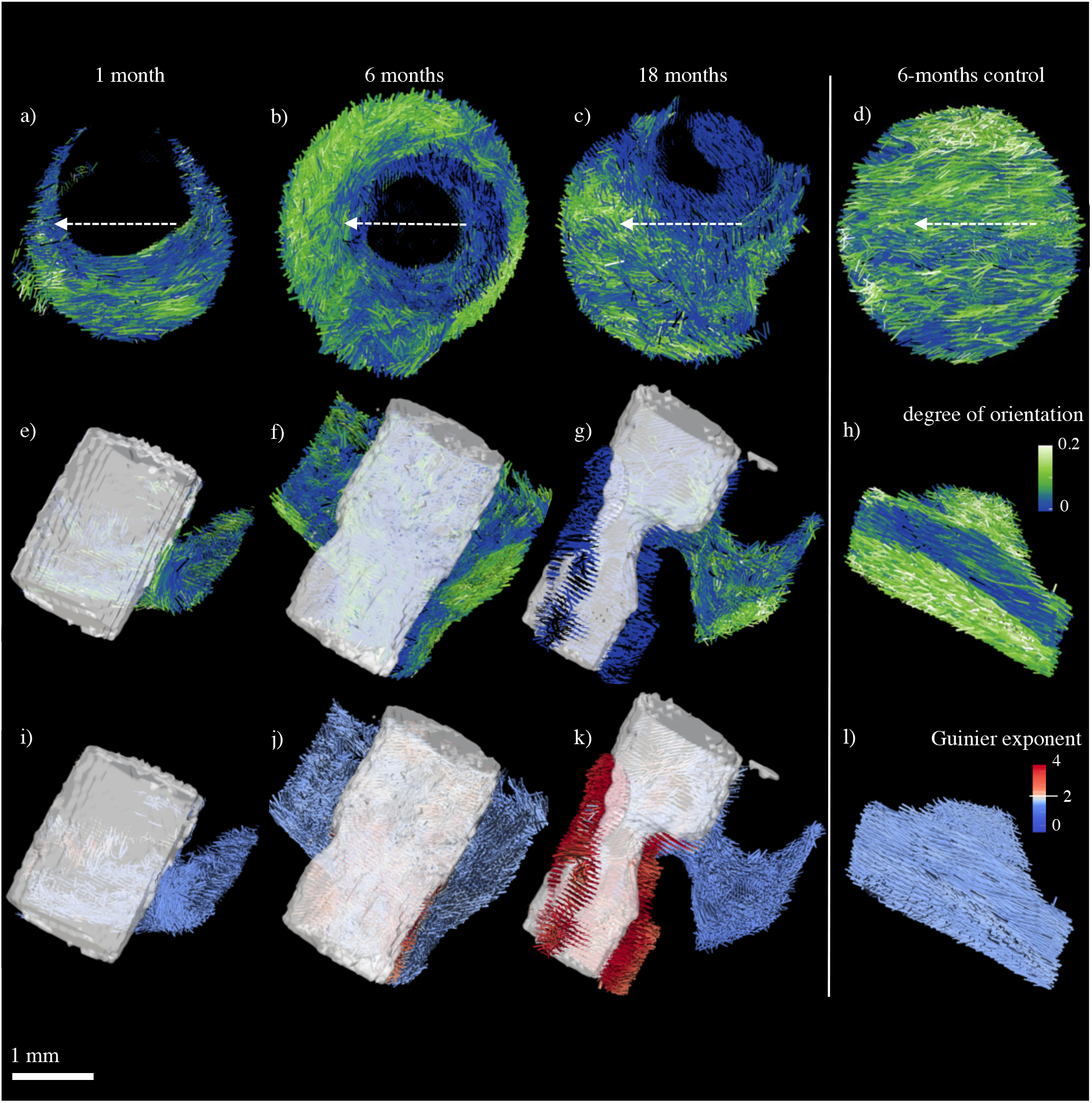
Reorientation of bone nanostructure represented by colored glyphs from SASTT reconstruction around a slowly degrading Mg implant (shown in white), from left to right 1 month, 6 months and 18 months after implantation plus a control sample 6 months after receiving sham treatment. The control sample after 1 month is presented in SI Fig. 2. The arrows in (a-d) depict the long-bone orientation from proximal to distal. The colors of the glyphs represent the degree of orientation (a-h) and the low-q exponent (i-l), respectively. The Guinier exponent shows higher values at the bone-implant interface. The glyph length correlates to the symmetric scattering intensity in all cases.

SASTT reconstructions in Figure 3 (a-h) depict the change in orientation and degree of orientation of mineral platelets in the vicinity of the degrading Mg implants. Figure 3 (a-d) reveals the view along the axis of the cylindrically shaped implant. The blank space in the middle corresponds to the implant locus, and the arrows indicate the longitudinal bone axis from proximal to distal side (see also Figure 2). Conversely, Figure 3 (e-l) shows the samples with a view perpendicular to the implant axis, where the remaining implant is displayed in white.

We observe after 1 month a very thin interfacial layer that formed around the implant (see Figure 3a and e). This layer is composed of a region in which the mineral particle orientation deviates from the expected global orientation along the bone axis and has a lower DOO. The generally low DOO of this sample is in part also due to the young age of the rat, and consequently ongoing bone growth, and compares well with the DOO found in the 1-month sham sample (Figure 3c, grey boxplot). The interfacial layer grows to about 0.5 mm with similar DOO after 6 months (see Figure 3b and f). In fact, nearly all of the measured volume of the 6-months sample shows an altered orientation, with the strongest effect visible at the interface. In contrast to this strongly altered orientation, the 6-month sham sample exhibits a rather strong orientation along the long-bone direction. The DOO is generally higher compared to the 6-month implant sample, and only shows some reduction towards the medullary cavity. After 18 months (Figure 3 c and g), a distinct preferred orientation is regained, and disordered zones are mostly found in areas where the implant is replaced by new bone.

In addition to the orientation and the DOO in a selected *q*-range, an extension of the data evaluation of SASTT was carried out to reconstruct the full scattering vector *q*-dependent intensity in every voxel, as described in the Methods section. This gives us sensitivity beyond the voxel size of the tomogram and allows us to obtain quantitative information in the SAXS size range, i.e. between 1 and 100 nm. These *q*-resolved reconstructions thus allow us to extract more detailed information on the size, shape and arrangement of the nanostructure. In order to do so, the average 3D directional scattering in each *q*-bin was considered from the full 3D reciprocal space map available. This generates a one-dimensional scattering curve of intensity vs. scattering vector *q* (Figure 4a), akin to standard SAXS experiments. This allows the determination of the thickness of the mineral particle via calculating the so-called T-parameter [7].

**Figure 4.**
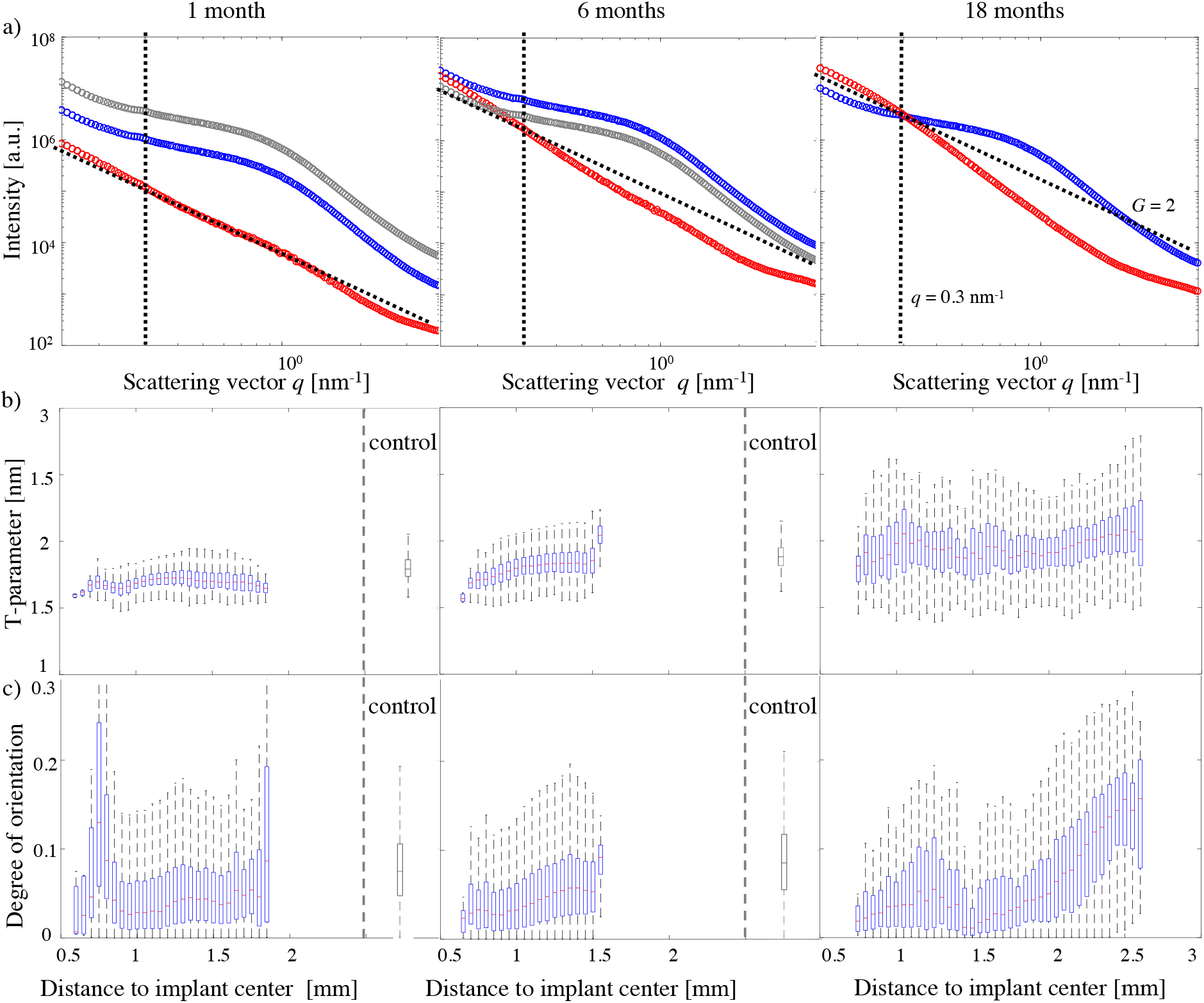
(a) Reconstructed scattering curve close to the bone-implant interface (red) and into the bulk of the bone (blue) at the three different timepoints (1, 6 and 18 months) and for the control (grey) at 1 and 6 months. (b,c) Boxplots of the T-parameter (b) and DOO (c), shown as a function of distance from the implant center. For 1 and 6 months, boxplots of the average T-parameter and DOO are also shown for the control samples.

Figure 4a shows scattering curves averaged within subvolumes from the interface region (red) and the bulk bone (blue) as well as from the control sample (grey). The shoulder formed by the scattered intensity as a function of *q* around 1 nm^-1^ is associated with the scattering of the mineral particles in bone. Here, we notice that this shoulder is mostly absent in the interfacial region. For all samples, the slope at low *q* (*q* < 0.3 nm^-1^) in the bone-implant interface region (red), referred here as *G*, markedly differs from that typically seen in mineralized bone. While in bulk bone the exponent *G* usually takes values between 1 and 2 [8], *G* values above 2 are found at the interface, as indicated in Figure 4a and Figure 3 i-l. In Figure 3 i-l the color code is set such that *G* values above 2 are represented by a red color. Comparing Figure 3i at 1 month after implant insertion with Figure 3j at 6 months and Figure 3k at 18 months reveals that the interface featuring higher *G* values has formed over time around the implant, in particular at sites with strong implant degradation and new bone formation. The control sample (Figure 3l and SI Figure 2) does not show any *G* values > 2. It is noteworthy that this very local change in the low-*q* exponent *G* does not fully correlate with a low DOO, which extends further into the bulk bone. Since the classical evaluation of the mineral particle thickness via the T-parameter is only valid for *G* ≤ 2 (as otherwise the determination of the Invariant from a Kratky plot is impossible), voxels with *G* > 2 (indicated in Figure 3 and Figure 4a) had to be excluded. Due to the finite *q* range accessible by the experiment, approximations have to be used for very small *q*, which become inaccurate at *G* > 2.

Figure 4 depicts the T-parameter (b) and the DOO (c) as function of implant distance; for details of the analysis see the Methods section. The box-plot representation chosen encodes statistical parameters such as the median (red line in the center of the box), interquartile range (size of the blue box), and the range of 1.5 times the interquartile range (dashed lines). A general increase in the T-parameter can be observed over time, which can be attributed to the growth of the animals and is also observed in the control samples (Figure 4). This, however, goes together with a local decrease at the interface for all timepoints and a spread in the interquartile range about 1 mm away from the implant center after 18 months. The DOO shows an increasing trend towards distances away from the implant surface. This is more pronounced over time and goes also along with a spread in the interquartile range over time. It is important to keep in mind that this data is obtained from one sample only due to the time constraints imposed by synchrotron experiments, in particular for the acquisition of the SASTT datasets. The analysis of whole 3D volumes instead of just 2D slices still produces detailed insight into the development of this sample set and we also ensured to choose representative samples in terms of implant placement and degradation state.

Figure 5 depicts the SEM analysis on cross-sections of the samples with a combination of back scatter electron (BSE) imaging (a) and energy dispersive X-ray analysis (EDX) (b-c). While the BSE is sensitive to the distribution of higher Z elements in the sample, the secondary electron (SE) signal is sensitive to the surface topology.

**Figure 5.**
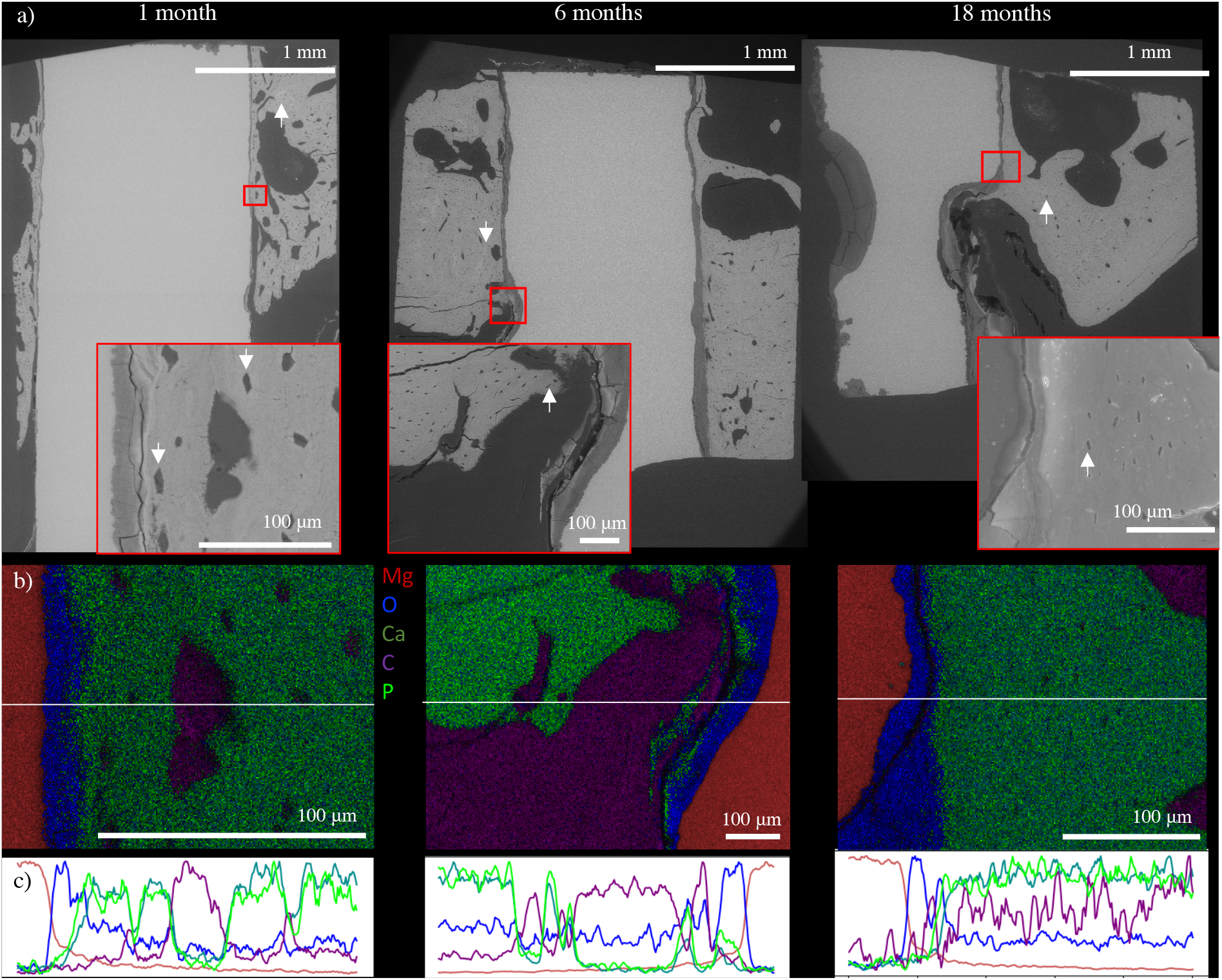
Scanning electron microscopy of cross-sections. (a) BSE overview images showing the morphology of the bone around the central implant. Lacunae are highlighted by arrows. Red squares mark the area of the magnified (in (b)) interfacial region of implant and bone. (b) Corresponding EDX maps, showing the elemental distribution at the bone-implant interface: Mg in red, O in blue, Ca in dark green, C in purple and P in light green. (c) Line profiles extracted along the white lines in (b), showing the interface layer as Mg/O-rich and revealing a gradual increase in Ca content towards the bone.

The imaging of the polished bone sections with BSE contrast is a well-established method to study mineralization processes [42]. While it is limited to 2D, it gives an overview of the whole bone cross-section sampled. The current approach allows us to visualize qualitative mineralization differences [43–45]. While the quantification necessitates for a proper calibration series [43], we use the BSE signal to qualitatively monitor mineral density differences in our samples, an approach already successfully applied to image mineralization processes in rat bone. [46, 47]

The BSE signal (Figure 5a) allows us to visualize the bone morphology and mineralization process by showing the lacunae (white arrows) and observing zones of varying mineral density (SI Figure 6, prominent zones marked with blue arrows). It is noteworthy that these zones are more prominent in the 1 and 6-months samples than in the 18-months sample. The overall lacunar density in the samples shows a slight decrease from 0.00049 lacunae/μm^2^ 1 month after surgery to 0.00042 and 0.00040 lacunae/μm^2^ after 6 months and 18 months, respectively.

Focusing on the implant-bone interface, an interface layer is visible which is co-localized with areas of an increased low-*q* exponent *G* (Figure 3i-l). The BSE signal indicates a lower Z composition than in the bone matrix, but still significantly higher than in the surrounding embedding medium. EDX was used in order to investigate the elemental composition of the interface layer. The regions investigated by EDX are highlighted in the overview Figure 5 (a) with red rectangles. The rationale for the selection of the various zones was to have implant-bone interface and embedding resin in the field of view. The slicing plane was chosen to align roughly with the virtual cut presented in Figure 3 (e-g). In Figure 5b, the different elements analyzed are represented in different colors: Mg in red, O in blue, Ca in dark green, C in purple and P in light green, which are overlaid to present their spatial localization. In the first step, the composition of the resin matrix and unaffected bone (as far from the implant as possible) was analyzed to generate a baseline for the identification of the elemental composition. The resin contains mostly C and O, whereas the bone is characterized by the presence of Ca and P. The EDX analysis of the interface region clearly allows to identify the Mg implant and the bone matrix from the strong signature of Mg and Ca, respectively (Figure 5b). The interface is characterized by a strong signature of O, very little Ca and a moderate amount of Mg. We thus conclude that this interface is composed mostly of Mg(OH)2, a commonly found degradation product of Mg implants [48, 49]. Its thickness varies between 50 and 200 μm for the different degradation states. Further away from this interface, a gradual increase in the Ca and P content can be observed, finally reaching a plateau value.

The SASTT measurements allowed for the first time to image in 3D the local misorientation of the mineral particles with respect to the normal direction in healthy bone along the main load direction, i.e. in the longitudinal direction of the bone shaft. For a quantitative description we used the normalized dot product between the local nanostructure orientation and the main load direction (reference vector, arrows in Figure 2 and 3). In addition, the normalized dot product between the local orientation and the implant orientation was calculated. This was done for every voxel and the corresponding results were averaged along the implant direction to generate a visual representation of the misorientation. A dot product of 1 indicates parallel orientation of the local orientation with the long axis of the femur (Figure 6a-c, g-h) or with the implant direction (Figure 6d-f, i-j), respectively, while a dot product of 0 conversely indicates perpendicular orientation.

**Figure 6.**
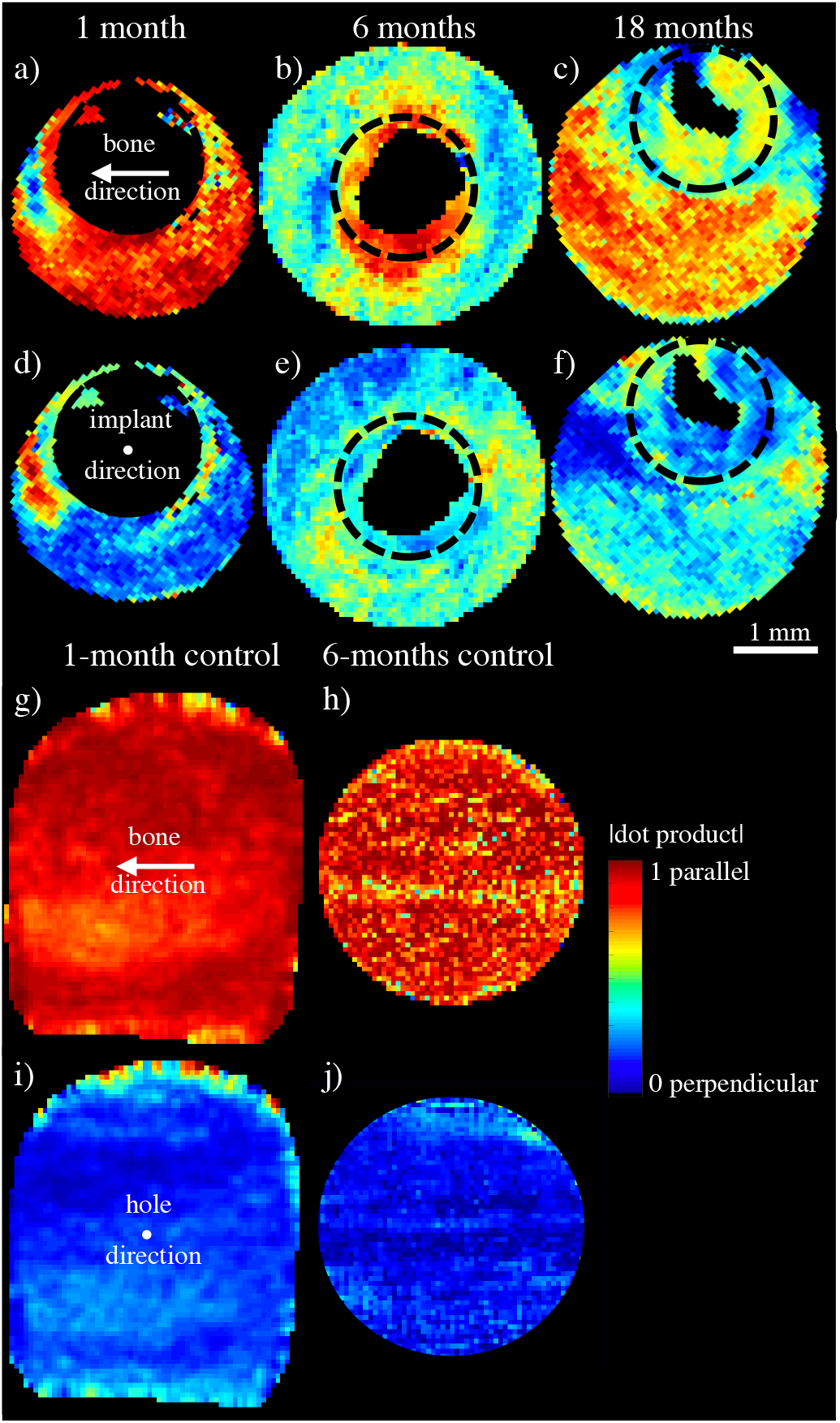
Dot product of the local mineralized collagen orientation with (a-c, g-h) the long axis of the femur (bone main axis) and (d-f) the implant direction or (i-j) hole direction for the control samples. The initial implant diameter is indicated by black circles.

The control samples show a parallel alignment of the mineral particles to the long-bone direction throughout the whole sample volume (Figure 6 g-h) without any re-orientation towards the hole direction (Figure 6 i-j). Figure 6a shows for the 1-month sample a preferred orientation of the bone nanostructure along the bone main axis, with strong deviations visible on the proximal and distal sides, where the misorientation of the local nanostructure with respect to the long axis of the bone is nearly orthogonal in a zone of approx. 400 μm. The comparison with the implant direction (Figure 6d) shows that the mineral-particle orientation, while being mostly orthogonal to the implant pin in the bulk of the bone, turns into an orientation along the implant direction close to the implant in the distal and proximal zones. After 6 months (Figure 6b), the general preferred orientation along the bone main axis is significantly reduced, and a strong reorientation around the implant can be observed. Especially in the distal and proximal zones, large areas of realignment can be found. In contrast to this, the compressive and tensile zones around the implant show a very strong alignment along the main bone axis, much stronger than in the surrounding bulk. The orientation with respect to the implant axis (Figure 6e) shows a more homogenous distribution, and in general the nanostructure shows a stronger alignment with the implant direction compared to the 1month timepoint. After 18 months, the alignment of mineral particles shows only a slight misorientation with respect to the bone main axis orientation in the distal and proximal zones (Figure 6c) and the bulk of the sample regained a strong preferred orientation. Also in the zones where the implant was replaced by bone, the nanostructure reveals a preferred orientation along the bone main axis. The comparison with the implant direction (Figure 6f) shows localized misalignment in the distal and proximal zones, while the bulk of the sample shows the same average orientation behaviour as the 6-months sample.

## 3. Discussion

We present the first study on 3D orientation of bone mineral particles around a degrading ZX10 Mg implant and show the subsequent nanostructural reorganization process involved during the healing period of 18 months. Previous studies, all performed via scanning in 2D, showed that the complex load situation in bone creates a nanostructural response in all three dimensions [50] depending on the orientation with respect to the anatomy, and in the case of implants on their type and implant placement [14]. Recent developments in analytical techniques strived to solve this problem by developing methods suitable for 3D investigations of nanostructured materials. First attempts relied on the investigation of sequential slices [51], whereas the development of tensor tomography used here allows for non-destructive investigations, permitting the study of 3D volumes for which consecutive microtome slicing is not possible without destroying the implant interface.

### 3.1. Growth and interfaces

The T-parameter extracted from the scattering curves is in the range of 1.5 to 2.5 nm, which is in good agreement with literature reports on healing bone [47], our own reports on Mg implants [13][12] and the control samples that were investigated in this study. In agreement with our previous studies, the T-parameter showed only mild changes, while other literature reports larger deviations during healing [52]. In this context it is, however, important to keep the deviations within individual samples in mind and the fact that the animals are still growing during the first six months of the study, which also leads to an increase in the T-parameter as visible in the control samples (Figure 4b). As the T-parameter assumes a degree of mineralization of 50% and the developing bone might deviate from this, corrections for this are employed in 2D studies. These rely on correlating the electron density derived from BSE with the T-parameter and calculating a corrected parameter, usually denoted as W-parameter [40] and defined as 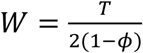, with *ϕ* the mineral fraction. As it is very difficult to obtain accurate mineralization densities for our 3D volumes, this correction could not be carried out in this case. As we mostly deal with lower mineralization levels in the early time points of this study, this lower mineralization will lead to a slight overestimation of the mineral particle size. Other factors to consider are the remodeling properties of rat bones, where the most notable difference between humans and rat bone is the absence of osteonal remodeling in rats and the associated usually low intracortical remodeling activity [53, 54]. Reports on the suitability of rats as an *in vivo* model conclude that, despite these shortcomings, the animals’ reactions towards the implant and the morphological response give valuable insights into the *in vivo* implant performance and the surrounding bone, respectively [55].

Through our BSE imaging, we are sensitive to mineralization differences and can thus follow the general bone development over time, displaying a gradual process from a woven-bone structure (1 month) to a more mature, laminar bone structure (18 months). Furthermore areas of active bone formation close to implant are still visible after 18 months (blue arrows in SI Figure 6). Globally, the lacunar density decreases with animal age, very similar to the process in human bone [56]. While lacunae are usually associated with osteocytic activity, SEM imaging cannot answer the question of whether they hosted viable osteocytes or not. The decreased lacunar density together with a more elongated shape and co-alignment of neighboring lacuna (Figure 5a) show a transition to a more mature bone [57], this is in line with the generally lower DOO of the young rat (1 month after surgery) compared to later time points (6 and 18 months after surgery) found with SASTT (Figure 2 and 3).

An interesting observation of this study is the development of an interface that grows and replaces the degrading implant, but does not exhibit the typical scattering of the bone mineral. This interface layer grows over time to a thickness of 50-200 μm. The scattering from this interface was found to exhibit a higher exponent in the small-*q* region, namely *G* > 2, which points towards a rather irregular structure. The interface is characterized by high levels of O and Mg, hence most probably a layer of Mg(OH)_2_, which has also been reported previously as a common degradation product [48, 49]. Further SEM investigations of the interface show a fibrous structure with granular patches (see red arrows in panel (c) and (d) of SI Figure 3-5). These zones, which exhibit a typical feature size of 300 nm, are particularly prominent in the 1- and 6-months samples. We hypothesize that these are Mg(OH)2 particles formed by implant corrosion. It is important to exclude these zones from the T-parameter analysis as it otherwise cannot be determined correctly.

At this point, we would like to emphasize that this kind of interface would be very hard to detect via scanning SAXS in a 2D slice, because the interface can be rather thin and we indeed were not able to resolve it in earlier studies [13]. However, other groups [16] identified a chemically different, amorphous calcium phosphate interface and ascribed its formation to the osteoinductive properties of Mg. It is in fact a strength of SASTT to give insights into complex arrangements, such as the pocket formation in the pits of degraded implant material, which subsequently mineralize.

### 3.2. Orientation

The orientation analysis allows to track the changes of the nanostructural orientation over time. Here, it is important to bear the mechanical load situation of bone in mind and how this influences the stress distribution around the implant. The mechanical optimization of bone is a research topic of great interest and there is general consensus that bone adapts to the current load situation and that drastic alterations in the load pattern (such as fractures) change the stress distribution and provoke a change in the nanostructure [47]. Re-orientation effects around implants have also been observed with SAXS [9]. In fact, it is well known that prolonged alterations in the stress distribution can lead to problems, where prominent examples refer to stress-shielding phenomena caused by a mechanical mismatch of the implant and bone moduli [58], albeit the moduli of Mg and bone are quite well matched [49]. While in this study it is hard to control the loading patterns, duration, or exact implant location, it is reasonable to assume a compressive/tensile zone from the animal anatomy, the walking activity of the animals, and the force due to gravity. Figure 2 illustrates a reasonable assumption for the predominant load situation in intact bone, whereas the presence of the implant and associated hole might, however, change this loading pattern.

In this study, we observe orientational differences between the mineral particles in the compressive/tensile zones and proximal/distal zones in the samples with implants, but not in the sham samples. We therefore interpret these findings as a distinct reaction of the bone to the presence of the implant, rather than a generic healing response to the insertion of a hole.

Our data did not support any significant differences between compressive and tensile or between proximal and distal zones. After the first month an initial reaction was observed, which was likely triggered by the modified loading situation and bone-healing processes, while implant degradation was still minimal. The nanostructure at the very interface points after one month along the implant direction (Figure 6a), forming a mineralized bridge between the cortical sides (Figure 6d), in accordance with previous investigations [13]. The main orientation along the bone axis, however, remains intact. As the implant degradation slowly starts and the animal also grows, an increased wrapping of the mineral particle direction around the implant can be observed after 6 months. This can be seen by the deviation from the main bone axis and extends over a large zone of about 1 mm (Figure 6b). In this case, the strong orientation differences orthogonal to the implant direction are reduced, while the overall level of the orientation along the implant decreases (Figure 6e). This may be interpreted as a mechanical adaptation by reducing the anisotropy in this direction. The zone affected might be larger than the sampled area, and other effects might be visible further away from the implant. In a previous study with a slightly larger sampled area we could, however, see that the impact of the implant placement on the nanostructural orientation extends to a zone of mostly 2 mm [13]. After 18 months, when significant implant degradation has already taken place, the nanostructure regained a strong preferred orientation along the main bone axis (Figure 6c), also visible in the regions where the bone had replaced the degrading implant. The images visualizing the alignment in the implant direction (Figure 6d-f) reveal that it mostly occurred in the zone around the implant, indicating a mechanical stimulus at play. It is worth noting that an alignment of the nanostructure parallel to the implant is particularly pronounced after 1 month, where the implant did not dissolve much, but the loading pattern changed due to the implant placement.

With the limited number of samples we can only speculate on possible mechanisms driving the nanoscale adaptation. Two major stimuli are identified, which are on the one hand the implant properties, such as the crystalline structure or the surface properties, and on the other hand the load conditions that are modified due to the placement of the implant and its subsequent degradation. While the chemical composition of a surface can have a large impact on the growth properties, this effect is expected to be most pronounced for the case of calcium phosphate surfaces [59]. A recent study using scanning SAXS could not show significant differences in the DOO or mineral particle thickness between different implant materials such as Ti, PEEK, or Mg-Gd alloys [60].

The second important factor to consider is the mechanical regime. It has been demonstrated by in-vitro experiments that osteoblast cells can align parallel to a surface or the direction of an applied mechanical stimulus, e.g. in response to cyclic stretching or fluid flow [61]. Aligned osteoblasts were shown to produce a highly orientated ECM with collagen fibers aligned in the same direction as the osteoblasts [62]. In bone, the situation is more complex, with osteocytes playing an important role as mechanoreceptors and likely also mediators for the 3D organization of the ECM [63, 64]. Mineralization of collagen fibers is known to be highly controlled by the collagen fibrils as a molecular template [65], resulting in elongated HAP particles aligned parallel to the collagen fiber axis and with the crystallographic c axis pointing in longitudinal direction as well.

We assume that the guidance by the implant surface and by mechanical stimuli is a major trigger of collagen/mineral reorientation observed in this study. An indicator for the importance of mechanical triggers in bone around implants was also found in a study investigating, among other parameters, the orientation of collagen/mineral around an implant in rabbit tibia by polarized microscopy [66]. The authors found that the orientation of collagen fibrils in the vicinity of the implant was altered when the implant was loaded cyclically. It should be noted, however, that a purely mechanical view might be too limited, since also wound healing processes take place. In healing bone disordered collagen fibers are typically deposited first (woven bone) which are later on replaced by orientated collagen fibers in a remodeling process. Due to the continuously changing healing front adjacent to a degrading implant and remodeling further away from it, these two processes are expected to happen in parallel. One can speculate that the initial response, the orientation of collagen fibers along the implant axis and formation of a bony bridge, is mainly governed by surface stresses induced by the implant. In the second stage the void created by the degrading implant has to be filled with bone tissue. The “wrapping pattern” in this stage has not been observed before, at least to our knowledge. It may have several reasons, such as a characteristic corrosion surface triggering the alignment of fibers in a circumferential way or local mechanical stresses that may be created by the corrosion process (gas formation) and potential inflammation. One can speculate that re-orientation in the direction of the main loading axis (longitudinal bone axis) starts only once these additional effects have subsided, therefore an intermediate stage might have to be considered which will also influence the mechanical stability during this time.

### 3.3. Limitations

This study was designed to investigate the impact of implant degradation and subsequent bone healing on the bone nanostructure, following up on previous studies by the authors. We have previously been able to follow the nanostructural rearrangement during progressing bone healing and implant degradation[13] and to characterize the impact of Mg on the bone nanostructure and mineral crystal structure. We also elucidated the bone-physiological response of fast Mg release[12]. However, several factors, such as the fast initial response towards implant placement, a possible out-of-plane wrapping around the implant and a hypothesized reorientation of the nanostructure during healing, remained unsolved questions in 2D and needed a 3D study to understand the sequence of orientational and nanostructural response. While the study presented here helps to understand how the degradation process affects bone nanostructure in 3D, there are also limitations.

Firstly, the number of samples is one per time point. This is owed to the limited access to synchrotron sources and the long measurement time (up to 24 h per sample) for this particular method. We tried to mitigate this effect by selecting representative samples based on CT scans of the explanted bones, excluding animals that showed any signs of abnormal implant degradation, and by ensuring proper placement of the implant during the initial surgery (see SI Figure 1).

Secondly, the chosen Mg implant pin in a transcortical geometry has limited medical relevance, contrary to choosing either an intramedullary nail geometry or a screw instead of a pin. The aim of this study was, however, to study the specific effect of the ZX10 Mg implant, where the pin geometry helps to reduce more extended geometrical factors as present in screws. Furthermore, the transcortical placement allowed us to investigate the influence of different loading situations in one sample, as opposed to intramedullary geometries. It is recognized that the geometry in this study is not directly applicable to the clinical case, but we are convinced that our key findings of altered mineral particle orientation will help guiding the development of next-generation materials for clinical applications and will thus facilitate the transfer to clinical studies.

## 4. Conclusion

In conclusion, this study sheds light on the 3D nanostructural adaptation of bone triggered by degrading ZX10 Mg implants. The main findings are the developement of the size of the mineral particles (T-parameter) induced by the implant degradation; the existence of a Mg-degradation product layer surrounding the degrading implant; and a sequence of two orientational motifs as a response to the implant placement and implant-degradation process. The first motif visible after 1 month is a strong orientation along the implant direction and a first slight wrapping of the nanostructure around the implant (Figure 6d), along with a weak reorientation along the bone main axis (Figure 6a). Subsequently, the second motif takes over, which reduces the orientation along the implant direction and induces a stronger wrapping of the bone nanostructure around the implant. Upon increased implant degradation, a normalization of the nanostructural orientation along the main bone axis can be observed, albeit an orientational signature along the previous implant direction also occurs (Figure 6c and f).

This study also shows that an extension of nanostructural studies towards X-ray scattering tensor tomography methods allows for new insights into bone-mineral restructuring upon implant degradation, such as the 3D orientational re-arrangement or the existence of thin interfacial layers around the implant, which would be difficult to draw from 2D scattering methods. In future research we will also exploit the fact that the reconstruction of the full 3D reciprocal-space map opens new possibilities for understanding the organization of nanostructures in 3D.

## Acknowledgements

Anders Palmquist is acknowledged for valuable discussion concerning the interpretation of the SEM data. The authors acknowledge the Paul Scherrer Institut for the allocation of two experimental sessions to acquire the data. X-ray experiments were carried out at the cSAXS beamline, Swiss Light Source, Paul Scherrer Institut, Switzerland. We would like to thank the Partnership for Soft Condensed Matter (PSCM) at the European Synchrotron Radiation Facility (ESRF) for support during the preparation of the experiment. The authors also acknowledge funding from the Area of Advance Materials Science at Chalmers University of Technology (ML), the Swiss National Science Foundation (SNF Sinergia, Grant No. CRSII5-180367) (JL), the Laura Bassi Center of Expertise BRIC (Bioresorbable Implants for Children, FFG – Austria) (EM, HE, AW), and the Berndorf Privatstiftung, Austria (HL, TG).

## Data availability

The authors declare that they have no known competing financial interests or personal relationships that could have appeared to influence the work reported in this paper.

## Data availability

The raw/processed data required to reproduce these findings cannot be shared at this time due to technical limitations (size of the datasets).

