## Supplementary Figures for "3D nanoscale analysis of bone healing around degrading Mg implants studied by X-ray scattering tensor tomography"

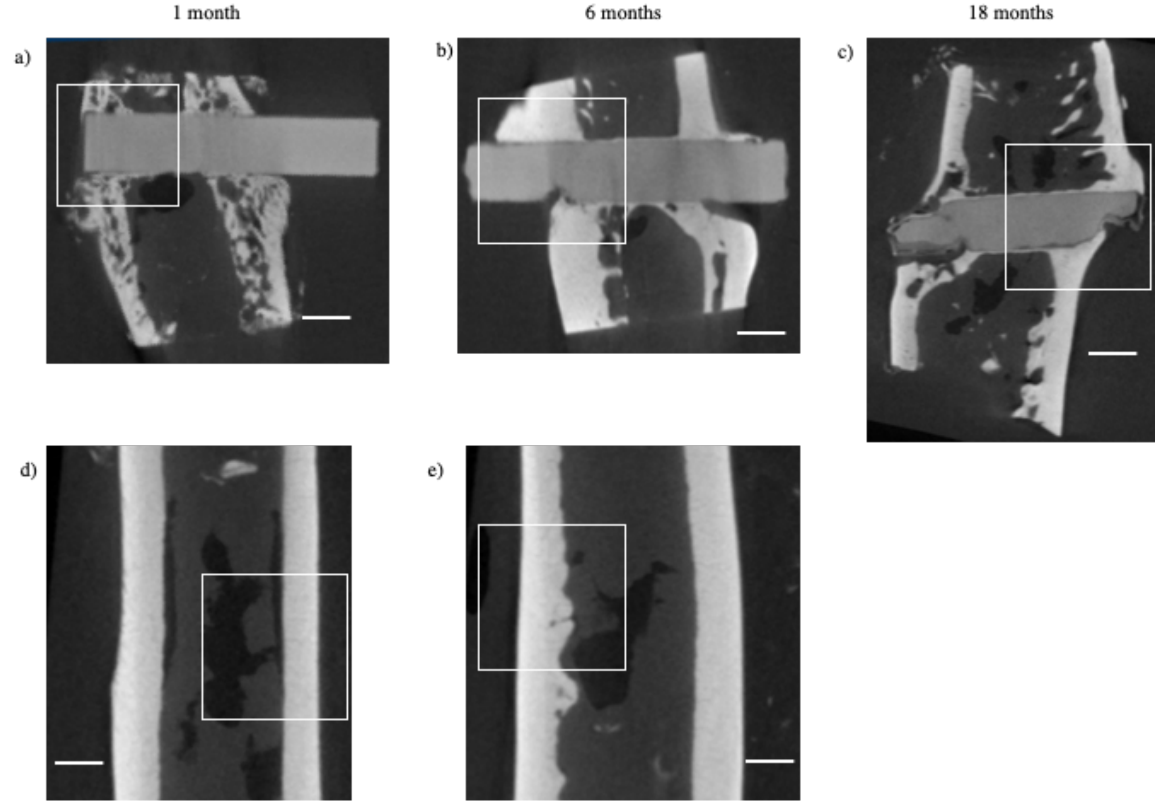


**Fig. S1.** µCT slices of the whole samples with sampled SASTT regions highlighted by a white square, for (a) ZX10 1 month, (b) ZX10 6 months, (c) ZX10 18 months, (d) sham 1 month, and (e) sham 6 months after implantation/sham intervention. The scale bar corresponds to 1 mm.

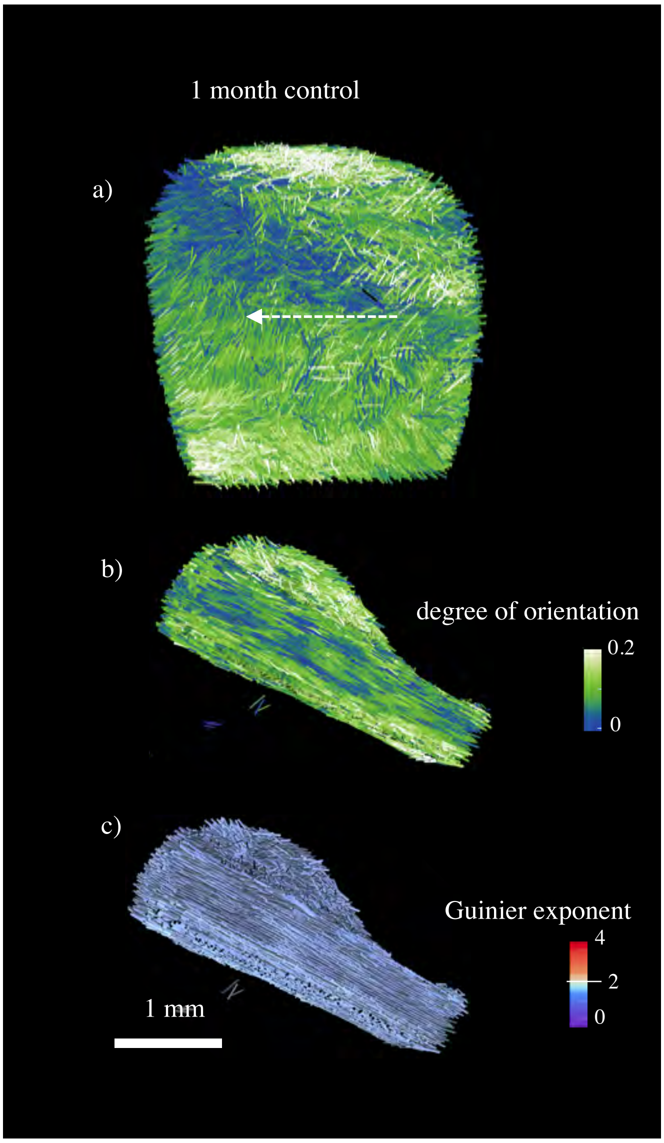


**Fig. S2.** Orientation of bone nanostructure represented by colored glyphs from SASTT reconstruction of a control sample 1 month after sham intervention. The arrow in (a) depicts the long-bone orientation from proximal to distal. The color of the glyphs represents the DOO (a,b) and the low-q exponent (c), respectively. The glyph length correlates to the symmetric scattering intensity in all cases.

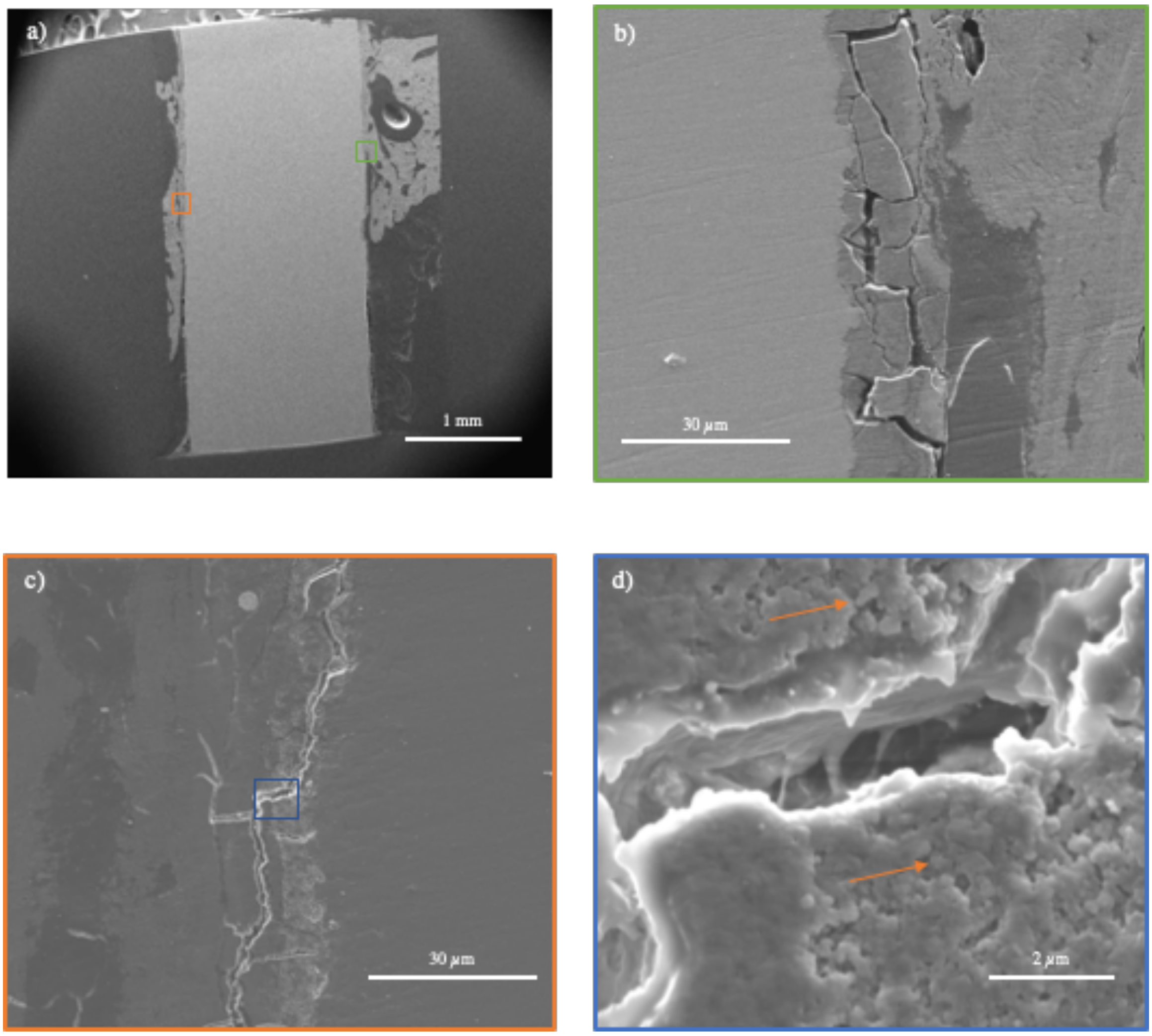


**Fig. S3.** SE SEM images of ZX10 1 month after implantation. The scale bars are indicated in the panels. (a) Overview of the whole sample with two ROIs indicated with green and orange rectangles. (b) Magnified green region. (c) Magnified red region, showing fibrous patches and granular matter. (d) Magnified fibrous patch region (blue rectangle in (c)), with granular matter highlighted by arrows.

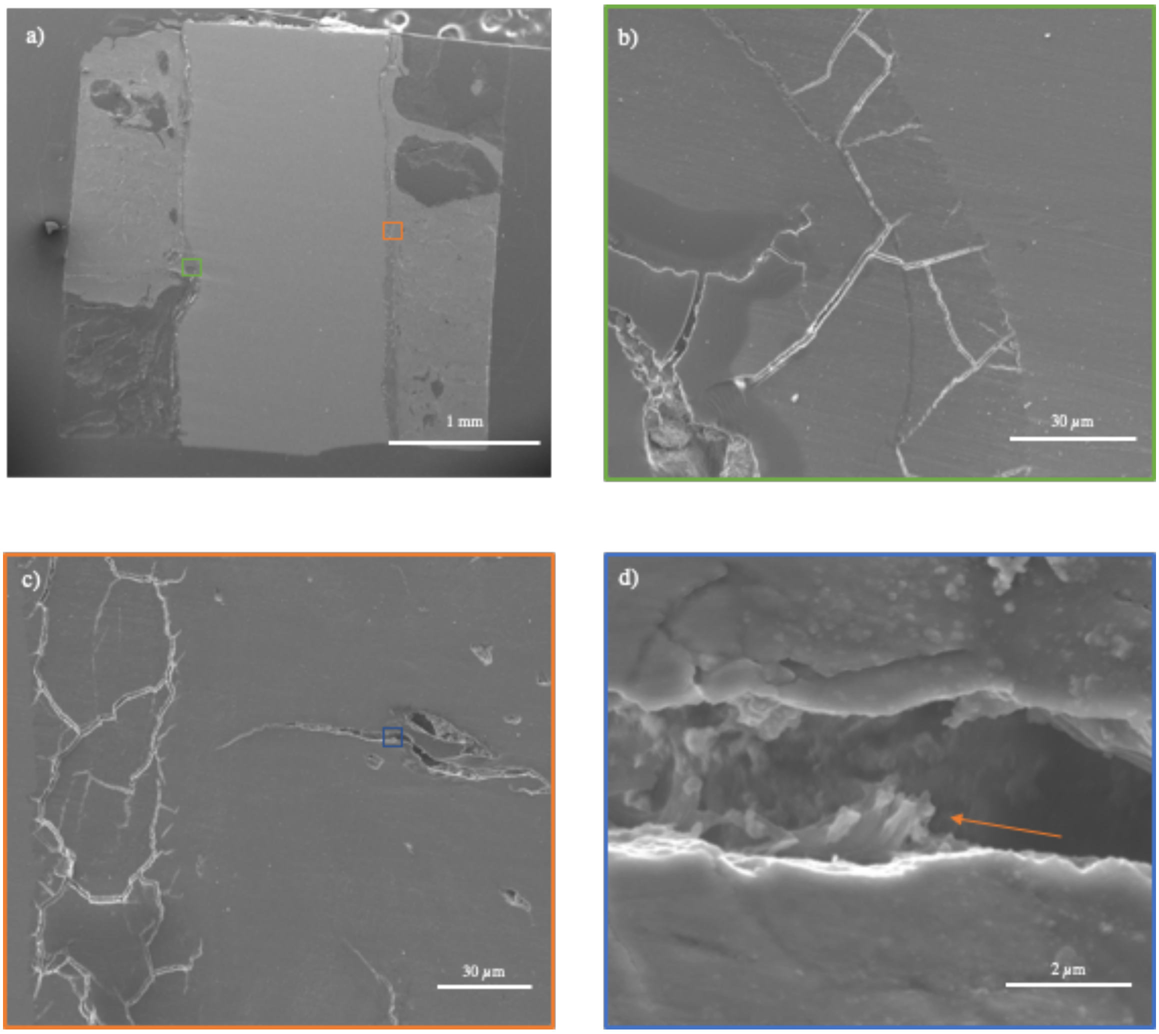


**Fig. S4.** SE SEM images of ZX10 6 months after implantation. The scale bars are indicated in the panels. (a) Overview of the whole sample with two ROIs indicated with green and orange rectangles. (b) Magnified green region. (c) Magnified red region, showing fibrous patches. (d) Magnified fibrous patch region (blue rectangle in (c)), with fibers highlighted by an arrow.

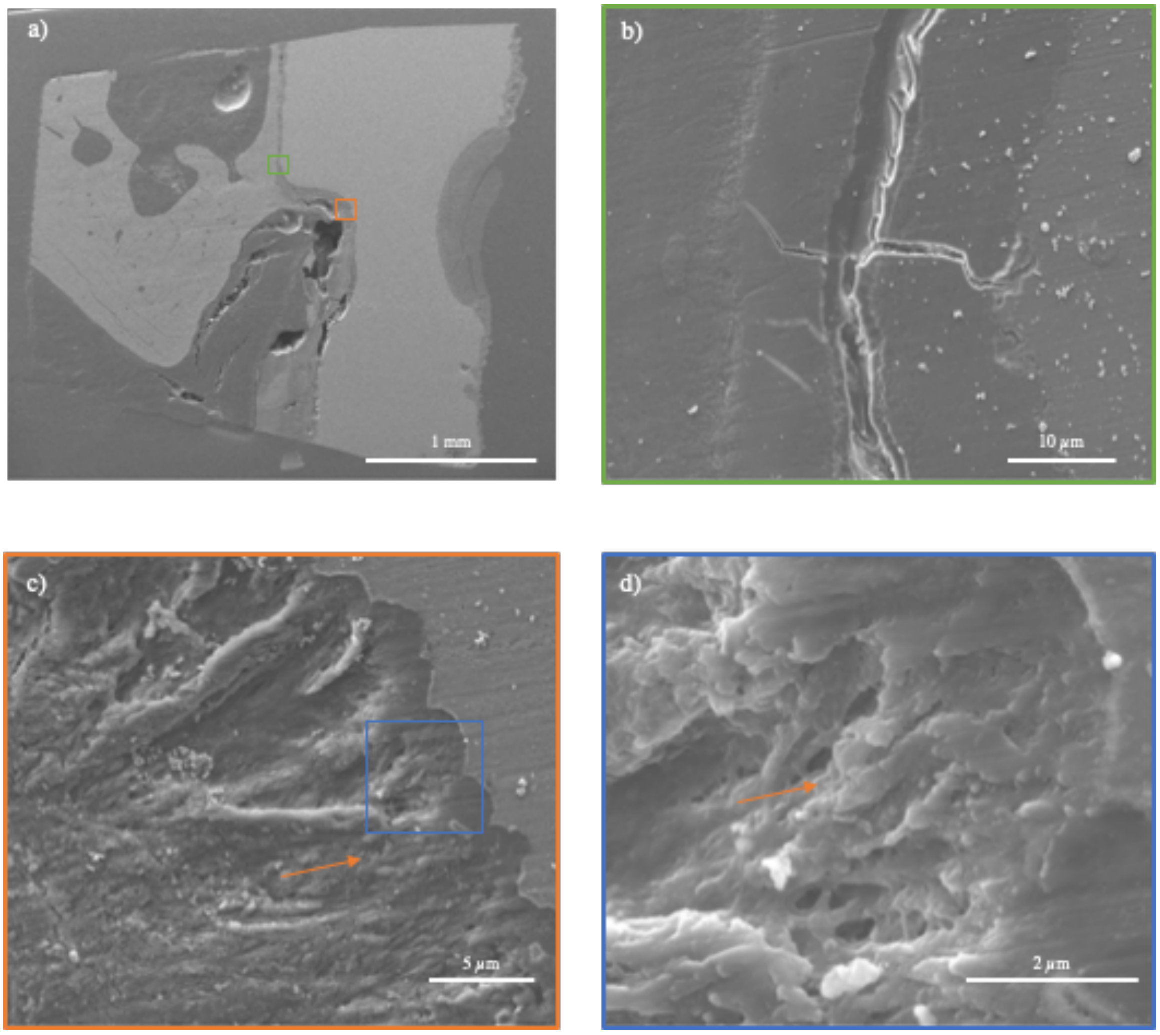


**Fig. S5.** SE SEM images of ZX10 18 months after implantation. The scale bars are indicated in the panels. (a) Overview of the whole sample with two ROIs indicated with green and orange rectangles. (b) Magnified green region. (c) Magnified red region, showing fibrous patches and granular matter in a strongly degraded implant interface. (d) Magnification of an exemplary fibrous patch region (blue rectangle) with granular matter highlighted by an arrow.

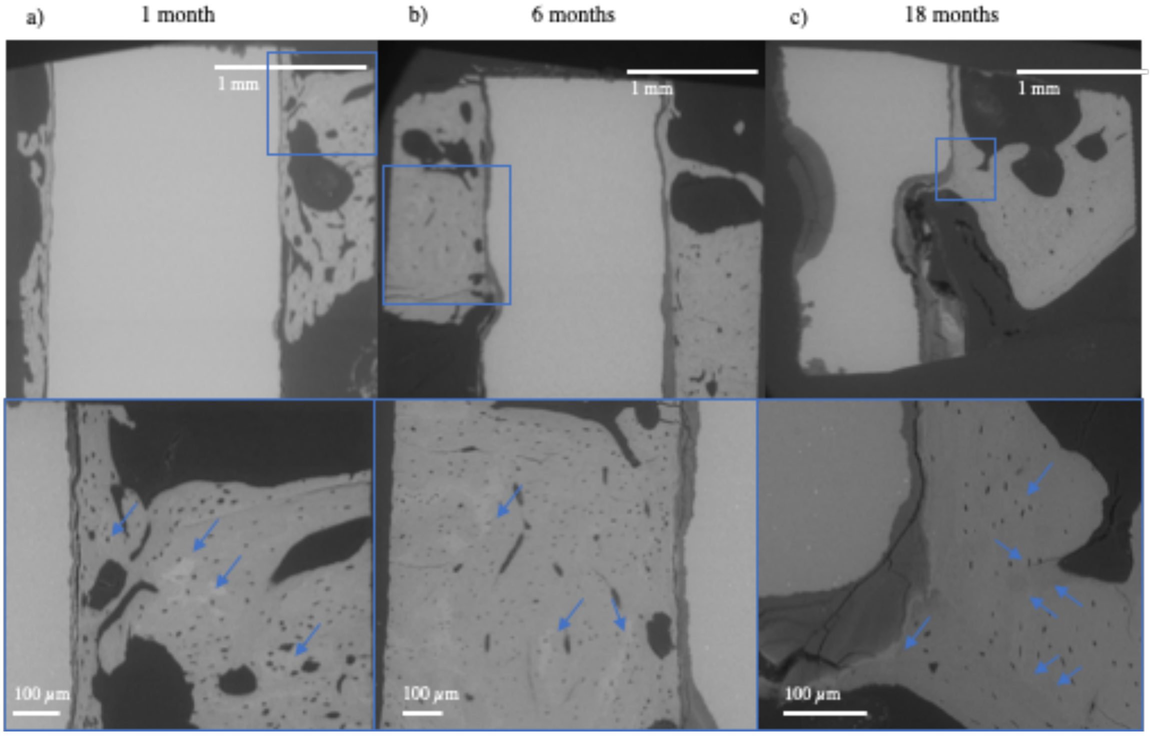


**Fig. S6.** BSE SEM images showing the whole sample (top panels) for (a) 1, (b) 6 and (c) 18 months after implantation. The bottom panels show magnified zones (blue rectangles), where distinct mineral density differences can be observed as indicated by the blue arrows.
